# Mechanosensation Mediates Long-Range Spatial Decision-Making in an Aneural Organism

**DOI:** 10.1101/2020.03.20.985523

**Authors:** Nirosha J. Murugan, Daniel H. Kaltman, Hong Jin, Melanie Chien, Ramses M. Flores, Cuong Q. Nguyen, Dmitry Tuzoff, Alexey Minabutdinov, Anna Kane, Richard Novak, Donald E. Ingber, Michael Levin

**Affiliations:** Department of Biology, Tufts University, Medford, MA, USA; Allen Discovery Center at Tufts University; Wyss Institute for Biologically Inspired Engineering, Harvard University, Boston, MA, USA; Steklov Institute of Mathematics in St. Petersburg, Russia; Department of Mathematics, National Research University Higher School of Economics, St. Petersburg, Russia; Harvard John A. Paulson School of Engineering and Applied Sciences, Harvard University, Cambridge, MA, USA; Vascular Biology Program and Department of Surgery, Boston Children’s Hospital and Harvard Medical School, Boston, MA, USA

**Author notes:** Author for correspondence: 200 Boston Ave., Suite 4600, Medford, MA 02155.

**Keywords:** *Physarum polycephalum*, unicellular, mechanosensing, basal cognition, decision-making, information processing, mass, stiffness, TRP channel

## Abstract

The unicellular protist *Physarum polycephalum* is an important emerging model for understanding how aneural organisms process information toward adaptive behavior. Here, we reveal that *Physarum* can use mechanosensation to reliably make decisions about distant objects its environment, preferentially growing in the direction of heavier, substrate-deforming but chemically-inert masses. This long-range mass-sensing is abolished by gentle rhythmic mechanical disruption, changing substrate stiffness, or addition of a mechanosensitive transient receptor potential channel inhibitor. Computational modeling revealed that *Physarum* may perform this calculation by sensing the fraction of its growth perimeter that is distorted above a threshold strain – a fundamentally novel method of mechanosensation. Together, these data identify a surprising behavioral preference relying on biomechanical features and not nutritional content, and characterize a new example of an aneural organism that exploits physics to make decisions about growth and form.

**Highlights:**

- The aneural Physarum makes behavioral decisions by control of its morphology
- It has a preference for larger masses, which it can detect at long range
- This effect is mediated by mechanosensing, not requiring chemical attractants
- Machine learning reveals that it surveys environment and makes decision in < 4 hours
- A biophysical model reveals how its pulsations enable long-distance mapping of environmental features

## Introduction

A defining feature of any living organism is its ability to select actions that maximize utility in diverse environments (Barandiaran and Moreno, 2006). Decision-making is a process that is well-characterized in behavioral studies of human and other vertebrate species possessing nervous systems. Increasingly, it is also recognized as a fundamental capability whose evolutionary roots extend to the most basal forms on the tree of life (Baluška and Levin, 2016; Lyon, 2006, 2015). Even unicellular organisms, including bacteria, have been studied as computational systems that collect information from their environment and use it to guide subsequent behaviors (Balazsi et al., 2011; Brunke and Hube, 2014; Danchin, 2009; Dexter et al., 2019; Norris et al., 2011). The emerging field of basal cognition focuses on the sensory, motor, and computational capacities of cells, and is thus tightly integrated with cognitive science, developmental biology, regenerative medicine, neuroscience, and even robotics (Baluška and Levin, 2016; Bongard et al., 2006; Doursat et al., 2012; Keijzer et al., 2013; Kriegman et al., 2017; Lyon, 2006; Slavkov et al., 2018).

Achieving a comprehensive understanding of the rules that underlie the related ways by which organisms make sense of their worlds will build a long-sought-after bridge between the biological and cognitive sciences. However, numerous knowledge gaps still exist. Understanding the abilities and limitations of decision-making in aneural systems is crucial for revealing the phylogenetic origin of complex cognition, and for the design of bio-inspired robotics and synthetic living machines (Ben Jacob et al., 2006; Ben-Jacob, 2009; Bronfman et al., 2016; Doursat and Sanchez, 2014; Doursat et al., 2013; Fields et al., 2019; Kamm and Bashir, 2014; Kamm et al., 2018; Keijzer et al., 2013). Moreover, it is an essential aspect of a biomedical roadmap seeking to identify stimuli for the rational control of swarm behavior of somatic cells during morphogenesis and remodeling, with many applications in birth defect repair, organ regeneration, and tumor normalization (Pezzulo and Levin, 2015, 2016). A central question for prediction and control of cell and tissue function concerns how evolution exploits physics for information processing and adaptive behavior in a wide range of materials and body architectures.

*Physarum polycephalum* is a remarkable organism that is being used to study decisionmaking and problem-solving in aneural biological systems (Vallverdu et al., 2018). Its basic structure consists of a series of concatenated tubules – a syncytium of nuclei and cytoskeleton that spreads out over centimeter to meter distances with the branching characteristics of vasculature. A unique feature of *Physarum* is their ability to coordinately shuttle protoplasm vigorously back and forth throughout their entire body in cycles with a regular period of approximately 2 minutes (Dietrich, 2015; Wohlfarth-Bottermann and Block, 1981). This movement, called shuttle streaming, allows the plasmodium to propel itself forward in any direction. *Physarum* is thought to drive shuttle streaming via contractile behavior that is effected by an intracellular network of cytoskeletal proteins comprised of both tubulin and actin subunits (Supplemental Figure 1) (Ueda et al., 1986). The slime mold uses this ambulatory ability to seek out nutrient sources and perform both chemo-and phototaxis (Durham and Ridgway, 1976; Häder and Schreckenbach, 1984; Natsume et al., 1993). As a polynucleated cell, the plasmodium also uses shuttle streaming to actively redistribute biochemicals and millions of nuclei which contain a genome consisting of 188 million nucleotides encoding ~34,000 genes (Alim et al., 2017; T.G., 1986), some of which contribute to signalling pathways that are crucial to multicellularity (Schaap et al., 2015).

*Physarum’s* deceivingly simple structure belies its complex problem-solving properties. Slime mold has become increasingly attractive as a model of proto-intelligence (Carello, 2012) and displays the major characteristics of a basic computational system that learns (Adamatzky, 2007; Kasai et al., 2015), including the capacity to identify the shortest path among many points (Reid, 2013), solving mazes (Nakagaki et al., 2000) and other spatial puzzles (Reid et al., 2012), habituating to noxious stimuli (Boisseau et al., 2016; Boussard et al., 2019), and anticipating periodic events (Saigusa et al., 2008). *P. polycephalum* has chemical -and light-sensing capabilities (Durham and Ridgway, 1976; Häder and Schreckenbach, 1984; Tanaka and al., 1987; Wohlfarth-Bottermann and Block, 1981). However, it is unknown what other physical mechanisms can be exploited by such a system. In complement to the existing work on local nutrient tropisms, here we sought to characterize how this organism might make decisions without prior exploration of a heterogeneous environment in the absence of light or nutrient gradients.

Here we present evidence suggesting that mechanosensation in *P. polycephalum* facilitates behavioral and growth decisions based on environmental features at long-range in the absence of nutrient gradients. We show that *P. polycephalum* can reliably detect and grow out towards different choices of chemically inert mass, and that this ability is abolished by mechanical disruptions such as tilting and mechanosensitive channel inhibition. We apply machine learning to reveal that information about the environment has been represented in the internal structure of the *Physarum* long before it takes morphogenetic action to directly interact with objects in its world, and propose a fluidically-coupled clutch-based model of long-distance information transfer to explain the experimental results. We find that *Physarum* displays an efficient goal-directed ability to make growth decisions by employing mechanosensation to collect information about its distant environment, revealing a novel behavior and mode for physics-based morphogenetic control not requiring a nervous system or multicellular architecture.

## Results

### *Physarum Polycephalum* makes weight-based decisions without environmental exploration

In the absence of chemical gradients or light stimuli, it is unknown whether or how *Physarum* engages with its environment or makes decisions about altering is morphology. In particular, it is unknown whether this architecture has any ability to sense features at a distance, without having to first encounter them directly.

To test the capacity of *Physarum* to make long-range decisions, we developed a discrimination assay to probe its ability to sense, discriminate, and determine outgrowth and motion between regions of differing mass in its environment. A piece of *Physarum* was placed in the center of a 10cm dish with two regions (at the dish’s edge) each containing a number of nonnutritive inert glass fiber discs. Each disc weighed 0.5mg (Fig 1A). Time-lapse imaging (Supplemental Figure 2) revealed that when presented with an environment containing three discs and one disc, the plasmodium preferentially grew towards the region of higher mass. Remarkably, this novel behavior was accomplished without first having to explore the area (Figure 1B). The first 12 hours of growth did not show any visually apparent directional preference, in fact the plasmodial front was equally distributed in all directions, indicative of the slime mold processing its environment. However, after 14 hours the plasmodium extended a branch directly toward the higher mass region. Thus, *Physarum* appears to have a preference for higher mass, which it can detect at a considerable distance from its body edge.

**Figure 1.**
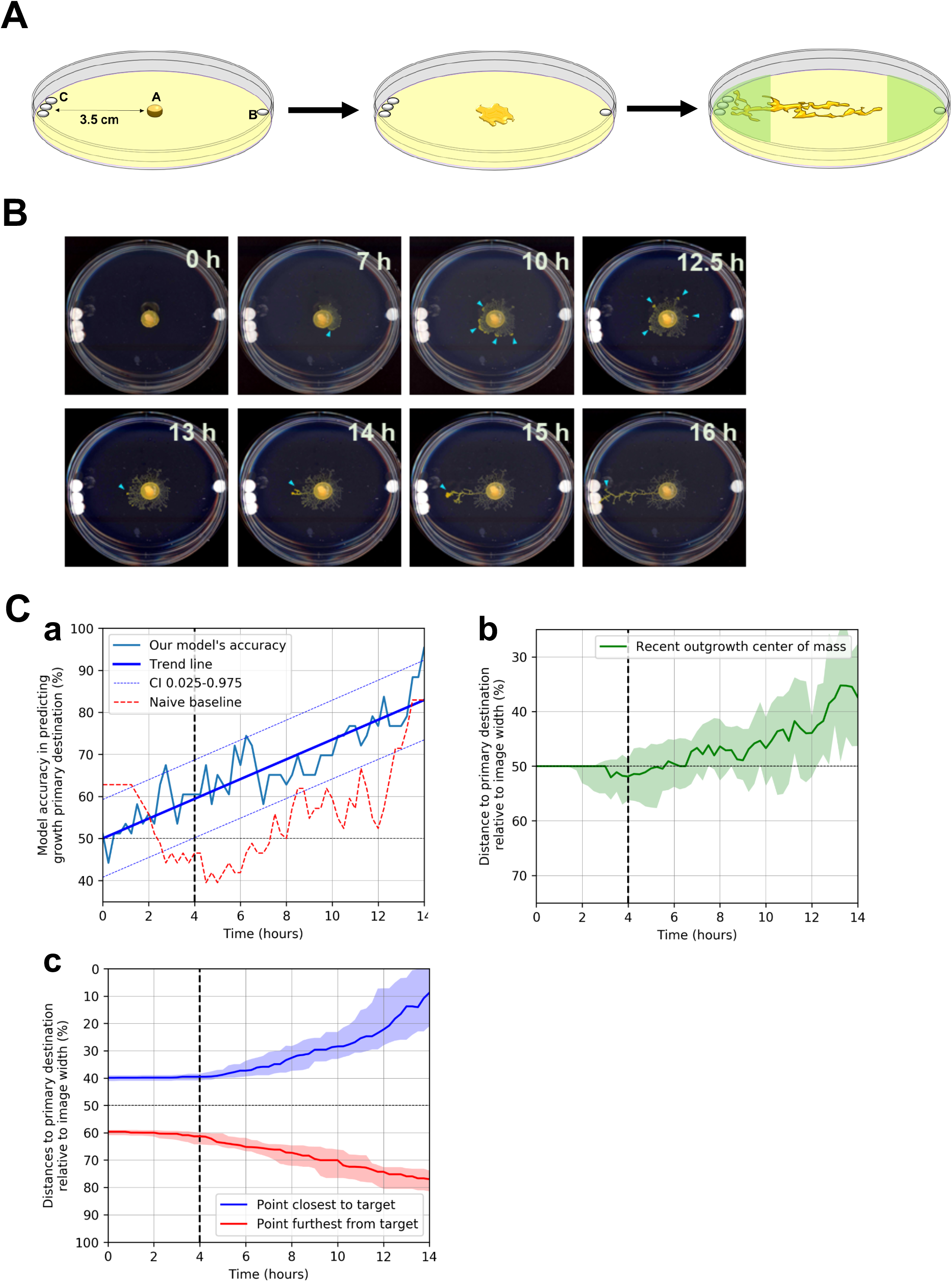
A novel mass discrimination assay revealed a preference for high mass over low mass in *Physarum*. (**A**) A mass discrimination assay was designed where inert glass-fiber discs were placed at the periphery of a petri dish coated with 1% w/v agar. In the most basic form of the assay, the *Physarum* (a) was plated between a low mass (b) region and a high mass (c) region containing 1 and 3 discs, respectively. (**B**) Time lapse reveals the incremental decision-making process. (**C)** Vertical dashed line indicates the time at which the prediction accuracy becomes statistically significantly higher than random, at this point the majority of the *Physarum* samples still grow in an opposite direction to their primary destination.(a) the light blue line corresponds to the mean accuracy for the prediction of the primary destination for each *Physarum* sample based on the observable history of growth; the thicker blue line corresponds to the trend in the accuracy improvement over time; thin dashed blue lines correspond to the 95% confidence interval for the measured accuracy; the dashed red line corresponds to the naive benchmark classifier that outputs the destination that the growth is closest to at the moment.(b) solid lines correspond to the median distances from the growth center of mass (green) and extremities (nearest — blue, furthest — red) to the primary destination; shaded areas correspond to the second and third quartiles of the population.(c) the solid line corresponds to the median distance from the recent outgrowth center of mass to the primary destination; shaded areas correspond to the second and third quartiles of the population.

We next sought to perform an objective, unbiased analysis of the growth process to better understand how early in its growth phase it had already made a decision. To determine the point at which directional information had become reliably available, we trained a dynamical machine learning model (Supplemental Figure 3) that was able to effectively predict the side the *Physarum* chose based on its historical behavior. Higher than chance prediction (59.4% accuracy, CI 95%:50.1-68.7%) could be made as quickly as 4 hours after being placed in the arena, long before actual outgrowth takes place (Fig 1Ca, dotted line). For a further 10 hours of imaging, the accuracy of the model increases by 0.59% every 15 minutes (Figure 1C). OLS regression for our model’s accuracy improvement over time yielded p<0.001, r^2^=0.946. Augmented Dickey-Fuller test rejected a spurious correlation. We also compared the accuracy of the machine learning model with the naïve deterministc classifier that simply outputs the side that the growth is closest to at the moment (Figure 1C, dotted red line). The naïve classifier performed significantly worse on the 3-13 hour interval, which concured with the observations by researchers. Early superior performance on the 0-2 hours interval prior to outgrowth was due to randomness in the initial shape that didn’t predict future outgrowth direction, which machine learning was able to learn well.

Interestingly, we found that in the first 6 hours of outgrowth the *Physarum* tended to slowly grow in a direction opposite to the target, afterwards accelerating to the target destination (Figure 1Cc, green line). The OLS regression model for the coordinate of the recent outgrowth’s center of mass confirmed this finding with p<0.0001. This unbiased technique revealed two phases of the decision-making process. In the first phase, sensing, which lasted as little as 4 hours, the *Physarum* had already detected the masses and decided towards which it will grow - analysis of its structure at that time could reveal an engram of the information it had gathered as a prediction of what the *Physarum* would do. In the subsequent phase, the *Physarum* executes the growth. Successful *Physarum* trajectories were also projected onto the axis spanning the two masses and plotted on a normalized time scale where 100% time indicates achieving the target (Fig. 1C). The results indicate a conserved process of minimal growth followed by a slight trend toward the smaller mass in the first half of the time frame. Only in the second half of the time frame is there consistent growth toward the larger mass. These findings suggest an initial measurement period of sensory integration and decision making followed by acting upon the decision by way of rapid outward growth toward the target.

### *Physarum* decision-making is constrained by distance and distribution of mass

Once the observable initial decision has been made, determined by observing a single outgrowth towards of the choices, it took only one hour for the plasmodium to cross the preestablished decision threshold and physically interface with the glass discs (Figure 2A). To statistically confirm this mass-based decision behaviour, we performed a χ-squared analysis that compared the observed and expected frequency of 4 possible decisions the organism could make: growth towards “low mass”, growth towards “high mass”, growth towards “both”, or indiscriminate growth, which was categorized as “none”. The expected frequency for the χ-squared test was determined by the *Physarum’s* inherent bias to grow in an empty arena (Supplemental Figure 4). When presented with the 3-discs versus 1-disc choice, the *Physarum* grew toward the 3 disc regions 70% of the time, while never choosing the 1 disc region alone. Interestingly, the proportion the plasmodium that selected either both regions or no region were only 12% and 18% (χ^2^=21.16, p<0.001, N=66) (Figure 2Ba) of trials respectively, suggesting that *Physarum* overwhelmingly made a choice when presented with regions of differential mass. Taken together, these results reveal that *Physarum* can detect differential mass at a precision of 0.1mg and preferentially grow toward the heavier mass.

**Figure 2.**
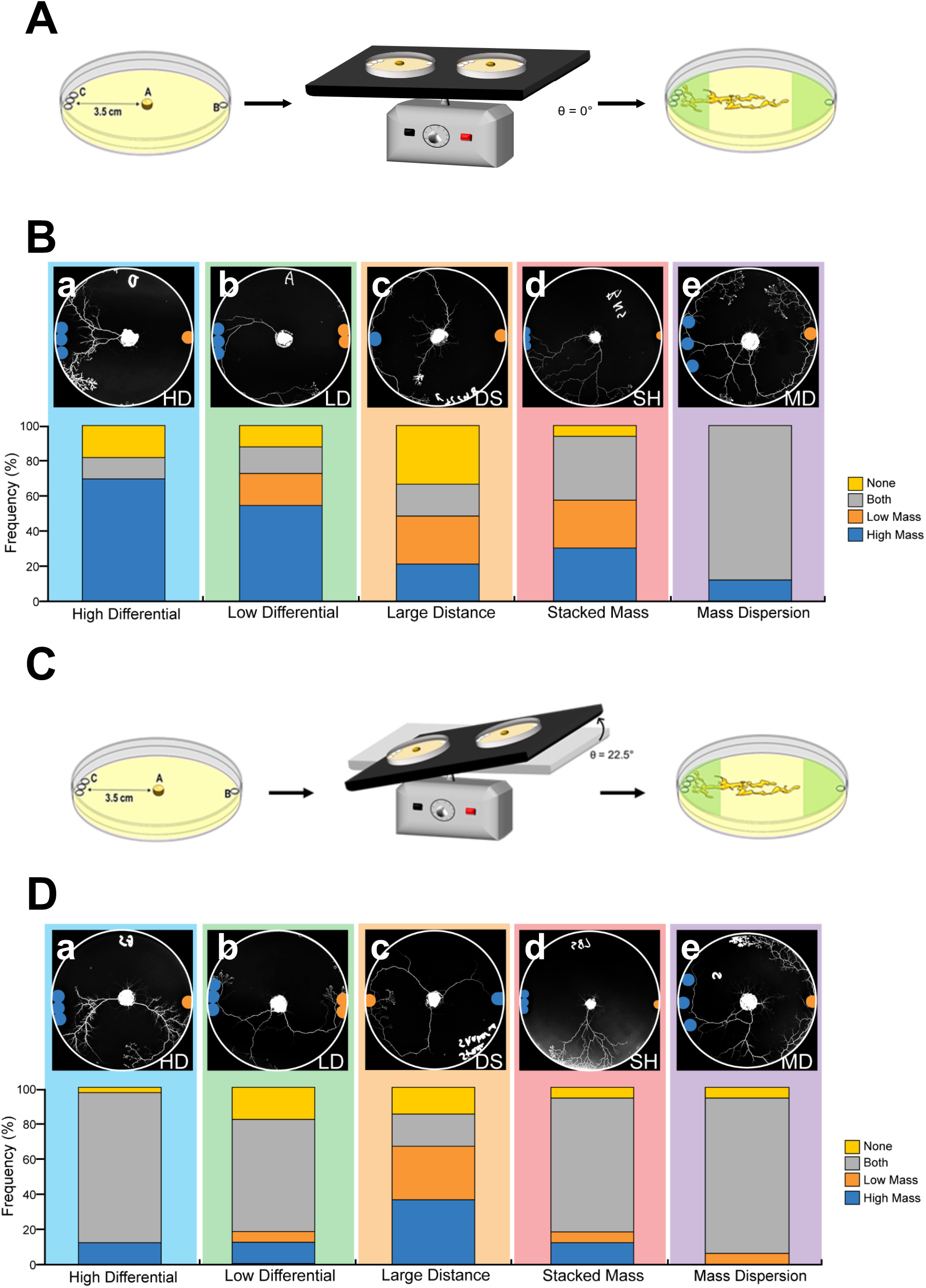
*Physarum* decision-making is constrained by distance, distribution of mass, and chronic mechanodisruption. **(A)** *Physarum* were grown in the mass discrimination assay on a flat, stable surface. (**B**) Mass differential, mass distribution, and arena size were systematically manipulated to investigate their influence upon mass sensing,.A χ–squared test was used to assess the difference between observed and expected frequency of 4 possible decisions the organism could make: growth towards “low mass”, growth towards “high mass”, growth towards “both”, or indiscriminate growth, which was categorized as “none”. The expected frequency for the χ–squared test was determined by the *Physarum’s* inherent bias to grow in an empty arena (Table 1). The *Physarum* was able to choose the higher mass in the (a) BA (χ^2^=21.16, p<0.001,n=66) and (b) LD (χ^2^ = 48.72, p<0.001, n=100) conditions, but could not discriminate in the (c) DS (χ^2^= 5.36, p=0.15, n=100) condition and preferentially chose both sides in the (d) SH (χ^2^=20.64, p<0.001, n=100) and (e) MD conditions(χ^2^= 48.72, p=0.001, n=100). Blue disc and orange disc are pseudocolored to show the assay more clearly. Significant differences between expected and observed outcomes within conditions are represented with *p<0.05. (**C**) The mass discrimination assay was modified by including mechanodisruption. *Physarum* were placed on the tilt table and continuously exposed to frequent movement. (**D**) Experiments from (**B**) were performed with continuous tilting to disrupt mass sensing. The vast majority (84%) of the *Physarum* selected both the 3-disc and 1-disc region (a) (χ^2^=30.15, p<0.001, N=100) or both the 3-disc and 2-disc region (b). (c) Tilt did not influence decision-making in large arenas (χ^2^=20.14, p<0.001, N=100). (d) In the stacked disc condition, tilt increased the number of trials where both regions were selected relative to the static condition (from 36% to 75%) (χ^2^=30.15, p<0.001, N=100). (e) When presented with dispersed discs in a tilted arena, *Physarum* continued to primarily select both regions (87% of trials), similar to the static surface assay (χ^2^=17.59, p<0.001, N=100).

**Table 1:**
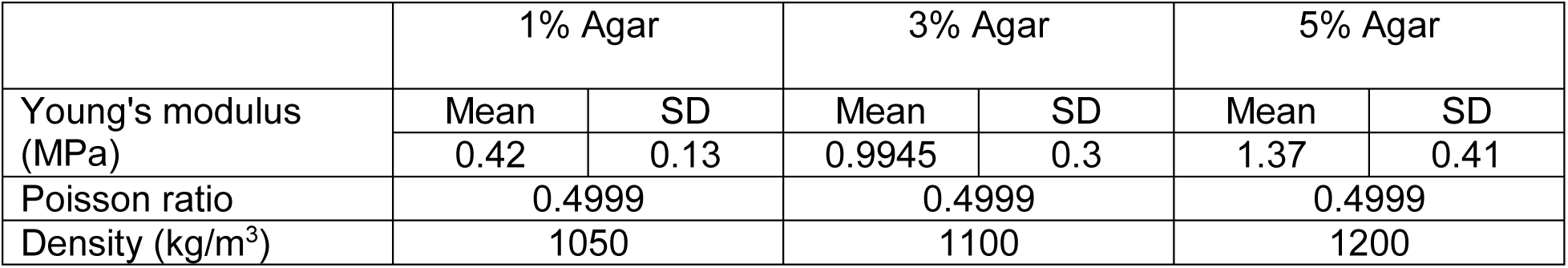
Parametric values for different agar concentrations

To characterize the limits of this mass-sensing ability, we modified our 3-discs vs. 1-disc base assay to include several variables that may impact decision-making outcomes, changing the mass differential, orientation, and inter-disc distance. When presented with a lower mass differential (3-disc vs. 2-disc), the *Physarum* continued to select the 3-disc region in 50% of trials, (χ^2^=48.72, p<0.001, N=100) (Figure 2Bb). The other possible options were selected in relatively equal proportions. When the *Physarum* was placed in larger arenas, decision-making was inhibited, suggesting that the distance at which decisions reliably be made is less than 25cm (χ^2^=5.36, p=0.15, N=100) (Figure 2Bc). When the area covered by the 3 discs was reduced by stacking, however, the *Physarum* was no longer able to distinguish between the 3 discs and the 1 disc, despite the mass differential. The “high mass”, “low mass”, “both”, and “none” options were associated with 30%, 27%, 36%, and 6% of cases respectively, (χ^2^= 20.64, p<0.001, N=100) (Figure 2Bd), indicating a 40% reduction in the ability of the *Physarum* to identify the region of higher mass. Spreading discs out laterally (increasing inter-disc space, Figure 2Be) produced a unique effect whereby the *Physarum* reliably selected both the 3-disc and 1-disc region in 88% of trials (χ^2^=48.72, p=0.001, N=100). This may reflect an inability of the *Physarum* to detect the 3 discs as one heavy individual mass and instead treat each disc as a separate entity of equal weight.

Based on these data, we conclude that *Physarum* can detect and direct its growth toward higher mass at a distance and is able to discriminate within 0.1mg at a distance of 3.5cm. Moreover, the propensity to choose a higher mass was decreased by lower mass differentials, reducing the area over which mass is distributed, and increasing the size of the arena. Interestingly, dispersing the high-mass region over a larger area made the *Physarum* consider both low and high mass regions as the same, growing towards both regions.

### Chronic mechanical disruption impedes mass discrimination by *Physarum polycephalum*

Because the discs were inert glass devoid of any nutrients and the assay was completed in the dark, the *Physarum* does not complete this task via chemical or light senses. Thus, we hypothesized that the *Physarum* utilized contractile activity to mechanically sense the physical aspects of its environment. In order to assess the importance of mechanosensation on long-range detection, we repeated the experiments described above (Figure 2B), but instead of growing the *Physarum* on a static surface, we chronically disrupted its environment by tilting the arenas on a rocker that subtended 22.5° from levelled surface at a frequency of 0.75Hz (Figure 2C). We compared the frequency distributions of plasmodia that were not tilted (Figure 2B) with those which were continuously tilted (Figure 2D).

We found that environmental disturbance impaired the ability of the *Physarum* to discriminate between low and high mass regions in every case. When making a decision in static environmental conditions, the high mass detection frequency was 70%, but when making the same decision during tilting, the heavy mass detection accuracy was reduced to only 10%. While tilting, the vast majority (84%) of the *Physarum* selected both the 3-disc and 1-disc region (χ^2^=30.15, p<0.001, N=100(Figure 2Da)). Similarly, tilted *Physarum* that were presented with a low mass differential (3-disc vs. 2-disc) test selected the high mass region in only 10% of trials relative to 50% when static (Figure 2Db). Whether *Physarum* were placed in small or large arenas, tilt did not influence decision-making as evidenced by a similar degree of non-specific outcomes (χ^2^=20.14, p<0.001, N=100. (Figure 2Dc). When discs were stacked in the high mass region, tilt increased the number of trials where both regions were selected relative to the static condition (from 36% to 75%) (χ^2^=30.15, p<0.001, N=100). (Figure 2Dd). Finally, when tilted and presented with dispersed discs, *Physarum* continued to primarily select both regions (87% of trials) similar to the static surface assay (χ^2^=17.59, p<0.001, N=100) (Figure 2De). In most cases, tilting *Physarum* resulted in a selection of both high and low mass regions rather than a preferential decision outcome. Together, these data suggest that exogenous mechanical perturbation prevented the *Physarum* from being able to discriminate between masses at a distance, consistent with the importance of the physical forces in the substrate and *Physarum’s* ability to detect these biophysical forces.

### Frequency-and inclination-dependant disruption of mechanosensation

As tilting *Physarum* was found to negatively impact mass sensing, we aimed to isolate the threshold at which environmental disturbance affected mechanosensation. First, we systematically decreased the frequency tilting with the aim of re-establishing high mass preference by removing mechano-disruptive interference. At 100% of maximum tilt frequency (RPM), *Physarum* preferentially selected both the 3-disc and 1-disc regions in 91% of trials, (χ^2^=15.76, p<0.001, N=95) (Figure 3A). Upon reducing the tilt frequency to 75% and 25% of the maximum RPM, both regions were selected in 71% and 30% of trials respectively (Figure 3A). Interestingly, as frequency of tilt decreased, high mass selection increased (Figure 3B). This clear shift from selecting both regions to selecting high mass regions with ever-decreasing tilt frequency illustrates the incremental negative contribution of mechano-disruption on *Physarum* mass sensing.

**Figure 3.**
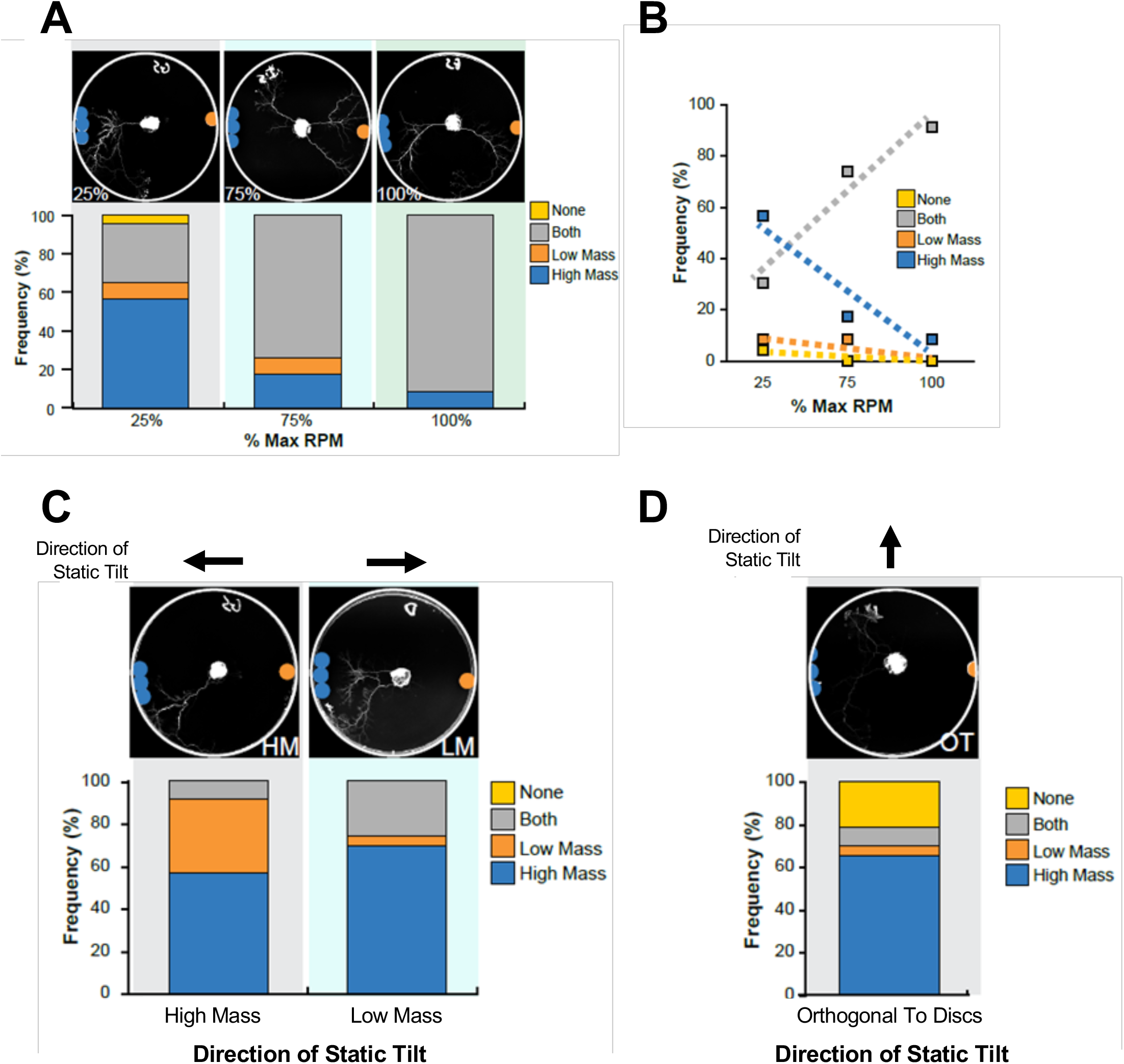
Slow rate of tilt and growth on an inclined surface do not affect *Physarum* decision making. (**A**) Disruption of mass sensing was frequency-dependent. As frequency of tilt increased (% Max RPM), high mass selection preference decreased. Instead, *Physarum* non-specifically selected both high and low mass regions with increased tilt frequency. (**B**)quantification of (**A**). (**C**) Plates were also placed on static platforms which were tilted to maximum inclination where the lowest point of inclination was directed toward the low or high mass region (black arrows), revealing increased high mass selection in the case of the latter. (**D**) Static tilting orthogonal to both high and low mass regions had no effect on *Physarum* decision making.

In order to ensure that the static physical incline was not preventing mass discrimination and that the perturbation of the mass-sensing was due to dynamic titling, we fixed the tilt table to its most extreme incline (22.5°) where either the 3-disc (high mass) or 1-disc (low mass) region was positioned at the lowest point of inclination. The table remained static in order to isolate the effects of incline alone from those of dynamic movement. When the high mass was positioned at the lowest point of inclination, the high mass region was selected in 70% of trials whereas the low mass region was selected in 4% of trials (Figure 3C), (χ^2^=11.43, p<0.001, N=95). When the low mass region was at the lowest point of inclination, the high mass region was selected in 56% of trials whereas the low mass region was selected in 35% of trials (Figure 3C) (χ^2^=9.77, p<0.001, N=95). When the lowest point of inclination was orthogonal to both the high and low mass regions, the high mass region remained the preferred selection in 65% of trials (Figure 3D) (χ^2^=13.01, p<0.001, N=95). While *Physarum* may prefer to grow in the direction of gravity, the incline itself does not prevent choice of either side. As the lowest point of inclination becomes increasingly biased toward the high mass region, high mass selection becomes more frequent. These data are consistent with mechanosensation that exploits a periodic event with specific frequencies.

### Orthogonal anisotropic gels weaken *Physarum’s* mass-sensing ability

We predicted, based on the hypothesis of mechanosensing of environmental features, that varying substrate stiffness would impact the *Physarum’s* capacity to rhythmically pull on the medium. Thus, we designed new arenas with anisotropic gel substrates composed of alternating lanes of poly-lactic acid (PLA) and 1% (w/v) agar (Figure 4A, CAD files in Supplemental Fig 5). Disc regions could be plated parallel to lane direction, on a common agar lane (i.e., 3discs-agar-Physar(agar)-agar-1disc) or orthogonal to lane direction, interrupted by lanes of alternating stiffness (i.e., 3discs-agar-PLA-*Physarum*(agar)-PLA-agar-1disc) (Figure 4B). As the *Physarum* grew outward, it was largely unaffected when growing parallel to the PLA lanes (Figure 4C) but became less able to deform its substrate when growing across lanes of PLA. Simulations of the experimental systems revealed dramatic differences in strain gradients between the two conditions (Fig. 4D), with the pattern orthogonal to the target direction resulting in a shortened propagation distance of strain (Fig. 4D inset). It was evident that when *Physarum* was plated on a common lane with both disc regions, high mass was preferred over other options in 60% of trials, (χ^2^=11.43, p<0.001, N=66) (Figure 4E). However, when plated orthogonal to lane direction, *Physarum* were unable to discriminate between regions and decision-making was inhibited (Figure 4F). The finding that an overly stiff substrate prevents appropriate mass detection suggests that *Physarum* is able to sense local strain to assess mass at a distance.

**Figure 4.**
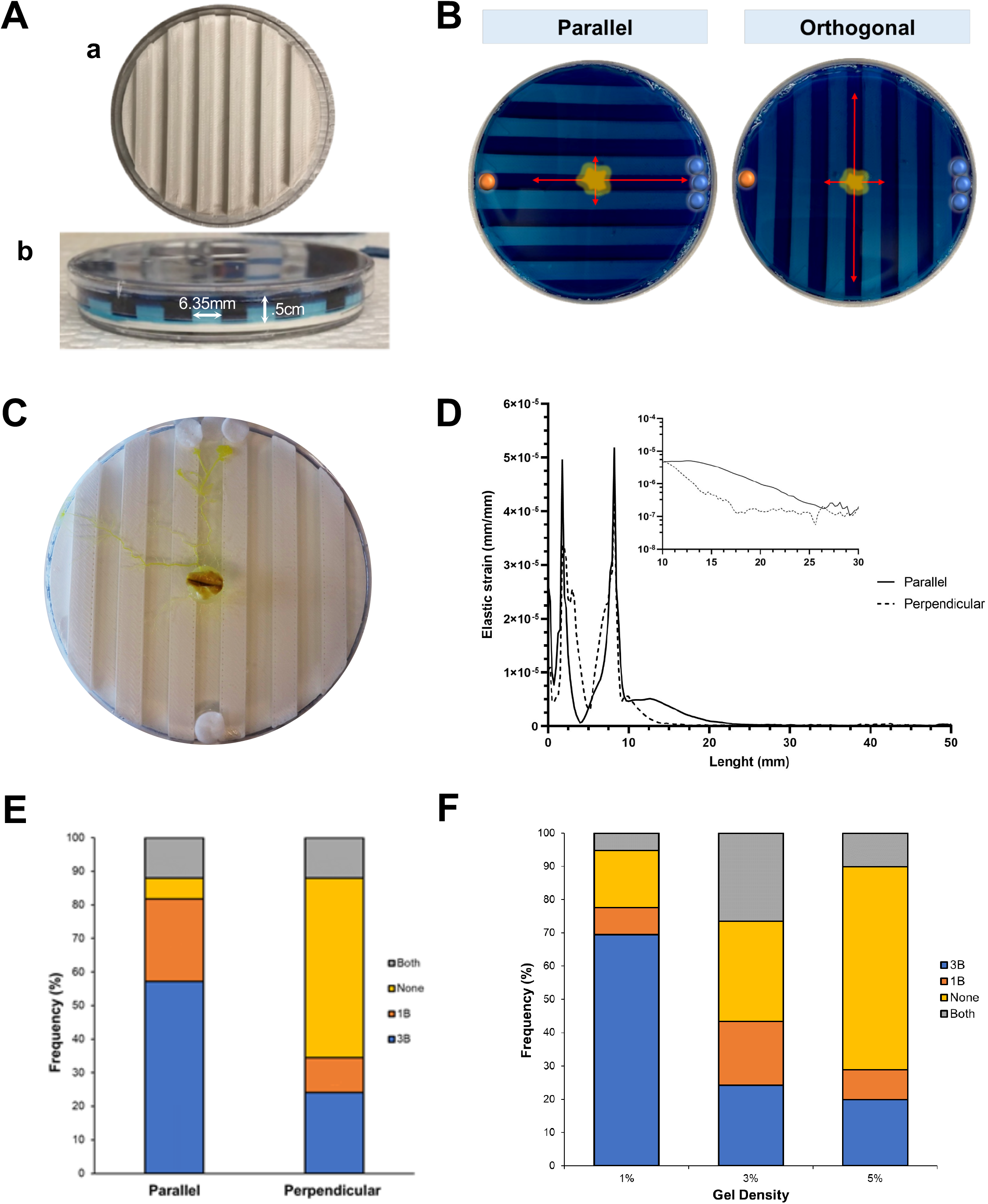
Anisotropic arenas were designed and built to assess the mechanosensing during mass-based decision making. (A) Arenas of anisotropic, alternating high and low stiffness substrate were fabricated using poly-lactic acid (PLA) molds (a). Agar (1% w/v) filled the wells in between each PLA lane, creating alternating 6.35mm high and low stiffness lanes (b, side view). (B) Discs were either placed on a common agar lane (i.e., disc-agar-*Physarum*(agar)-agar-disc), parallel to all other lanes or placed orthogonal to lane direction with alternating high and low stiffness regions distributed between discs (i.e., disc-agar-PLA-*Physarum*(agar)-PLA-agardisc). (C) *Physarum* growing on an anisotropic gel where the assay was run parallel to the plating. The perpendicular condition had discs plated 90° off of current placement. (D) Plot of agar strain along the centerline from the edge of the plate at the target mass toward the center of the plate showing much reduced strain propagation in anisotropic gels oriented perpendicular to the strain gradient. (E) When plated on a common agar lane, parallel to all other lanes, *Physarum* preferentially selected the high mass region; however, when plated orthogonally, *Physarum* was non-selective. (F) Arenas were then fabricated to test the effect of gel density on decision-making. Low-density gels (1%) were associated with high mass selection whereas increased densities (3 or 5%) contributed to decreased selectivity.

As stiffness likely impacted the strain fields across the agar substrate, we fabricated new arenas that varied by gel density (Figure 4E). The *Physarum* was plated on a 1%, 3%, or 5% agar arena and subjected to the classic high-low mass paradigm (i.e., 3-disc vs. 1-disc). We observed that low stiffness (1% agar) arenas were most conducive to high mass selection, with *Physarum* selecting the 3-disc region in 70% of trials, (insert chi-squared statement) (Figure 4E). Higher stiffness conditions did not generate similar levels of discrimination and higher stiffness was associated with more non-selection (Figure 4E). Taken together, these data demonstrate that both the distribution and magnitude of substrate stiffness can impact *Physarum*’s mass sensing ability.

### Long-range Mass Sensing is TRP-channel dependent

Given the known pulsing activity of *Physarum* (Coggin, 1996; Dietrich, 2015), we hypothesized that the mechanism by which the plasmodia senses mass involves mechanosensation that is mediated through shuttle streaming and contraction, in which the *Physarum* rhythmically pulls on the medium and interprets the traction force information that is altered by objects in its vicinity. This kind of force sensing is known to be mediated by stretch-sensitive ion channels in several other systems. Thus, we hypothesized that the ability to direct initial growth toward heavier mass would require the function of stretch-sensitive channels such as TRP-like channels. We used the blocking peptide GsMTx-4, which has been shown to be a potent TRP channel inhibitor (Gnanasambandam et al., 2017). This peptide is predicted to block a number of TRPC-like proteins in the *Physarum* genome (Glockner et al., 2008). In these assays, 30uM GsMTx-4 was pipetted onto the *Physarum* 30 minutes prior to the assay to ensure the drug would be fully absorbed prior to the mass-sensing assay was carried out. The *Physarum* was then allowed to make a decision in our base assay under the influence of this TRP-channel inhibitor. We observed a statistically significant difference between expected and observed frequencies of decision categories when presented with a binary choice of a 3-disc (high mass) and 1-disc (low mass) region (Figure 5A), χ^2^=52.58, p<0.001, N= 90), indicating that under the influence of the TRP channel inhibitor, *Physarum* only selected the high mass region in 11% of the trials, while selecting both high and low mass regions in 71% of trials (Figure 5B). In contrast, plasmodia exposed to vehicle (water) selected the 3-disc region in 87% of trials; the “low mass”, “both”, and “none” categories were selected in 4%, 7%, and 2% of trials respectively (Figure 5B). When *Physarum* was exposed to NIR001, a linearized peptide control for GsMTx-4, the high mass region was selected preferentially in 65% of trials, (x^2^=22.44, p<0.001, N=90). Together, these data suggest that mechanosensitive TRP channels are required for *Physarum* to make massbased decisions.

**Figure 5.**
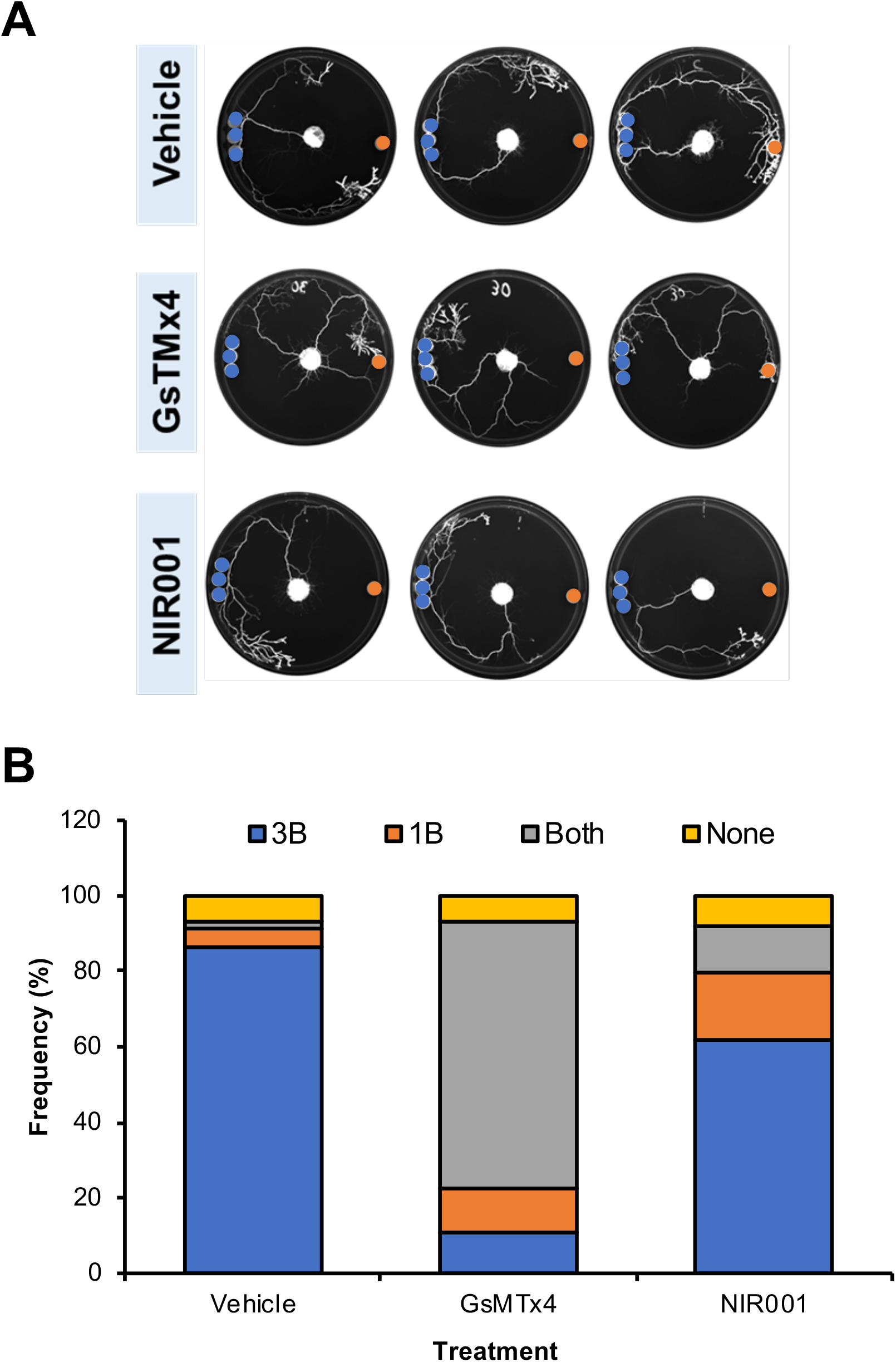
GsMTx-4 treatment disrupts mechanosensation in *Physarum*. (A) Representative mass discrimination trials are displayed for *Physarum* treated with vehicle, GsTMx4 (a TRP channel-blocking peptide), or NIR001 (a linearized peptide control for GsTMx4). (B) *Physarum* was able to select the high mass regions upon exposure to vehicle or NIR001, but GsMTx4-exposed *Physarum* non-specifically selected both high and low mass regions.

### Dynamic Compression device-induced pressure waves disrupt mass sensing

As we suspected that strain fields were the operant variables underlying mass sensing, we asked whether temporally changing the fields might affect decision-making. In all other experiments, static mass conditions were utilized, and it was consistently revealed that high mass regions were generally preferred unless a mechano-disruptive variable was introduced. We therefore hypothesized that dynamic compression of the substrate would influence *Physarum* decision-making. To test the hypothesis, we replaced the high mass region with a moving piston and pulsed the substrate with different patterns of compression; we continued to place 1 glassfiber disc at the opposite side of the arena (Figure 6A). When pulsing was matched to the *Physarum’s* average flow rate, the pulsed region was overwhelmingly preferred and was selected in 90% of trials, (x^2^=19.74, p<0.001, N=66) (Figure 6B). Pulsing the substrate to match *Physarum* shuttle streaming or as a non-specific sine wave produced pulse preference in 50% and 40% of trials respectively (Figure 6B). Indeed, when matching the pulse pattern of substrate compression to the average *Physarum* flow rate, the 1-disc region was never selected over the pulsing region (Figure 6B). The loss of preference for the pulsing side when the pulsing matches shuttle streaming provides further evidence that the shuttle streaming activity is what drives mass sensing at long distances.

**Figure 6.**
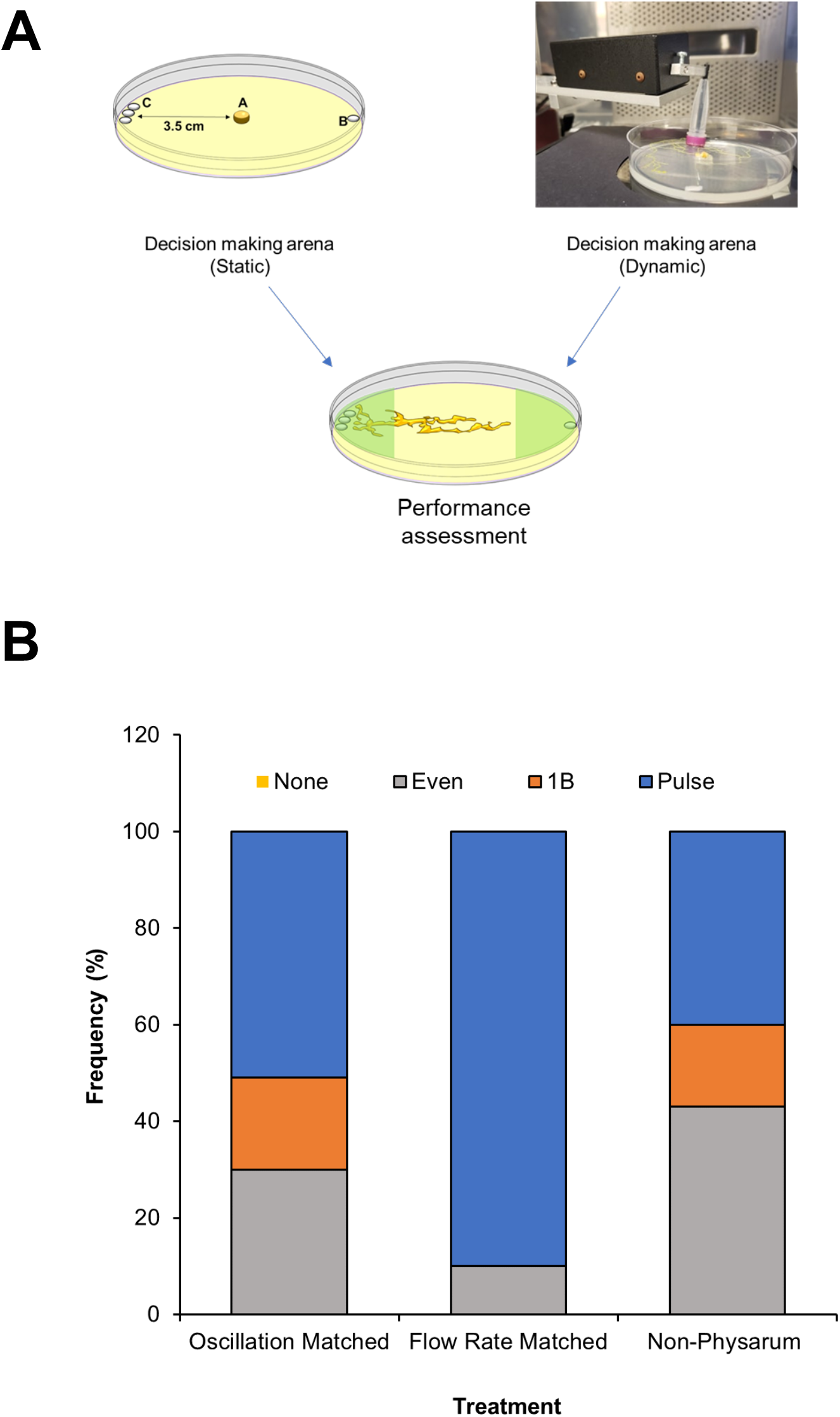
Dynamic pulsing of the agar substrate impacts decision-making and is pattern dependent. (A) In addition to static disruption of strain fields presented elsewhere, we constructed a dynamic disruption paradigm. *Physarum* were plated in the center of a standard arena; however, the target regions consisted of either a 1-disc (low mass) condition or a piston that periodically compressed the agar substrate. The pattern of compression could be presented as a non-specific sine wave or controlled to match *Physarum’s* average oscillation (0.01 Hz), or flow rate. (B) Flow rate-matched pulsing was overwhelmingly preferred relative to other options; All conditions, including oscillation-matched pulsing and non-specific sine waves were preferred over 1-disc regions.

### Computer simulations of agar stress and strain

To gain greater insight into the physical cues that *Physarum* senses using this mechanotransduction mechanism, we carried out finite element modeling of stress and strain profiles in the culture substrates. As expected, larger masses increased the magnitude and width of strain gradients (Figure 7A) at a radius of 3cm from the center, indicating that if *Physarum* can detect strain field gradients it could be sufficient to facilitate the choice of larger masses over smaller masses provided sufficient sensory precision (Figures 1-3). Additionally, the greater stiffness of the higher percentage agar reduced the magnitude and width of strain gradients (Figure 7B), reducing the ability of the *Physarum* to detect mass at a distance, and conversely, lower percentage agar increased the magnitude and width of the strain gradients (Figure 7B). These predictions were supported by experimental results showing that the ability of the *Physarum* to detect mass at a distance dramatically reduces with increasing agar percentage (Figure 4F). Finally, the finite element model was able to explain the unexpected observation that the *Physarum* could not distinguish between 3 stacked discs and 1 disc, but could distinguish between 3 discs placed next to each other and 1 disc. In order to compare strain profiles in our simulations, we assessed the Threshold Strain Horizon Angle, the angle that encompasses the region where strain is above the minimal strain sensitivity (1.49×10^-7^), the strain on 5% agar at which decision-making performance breaks down. The simulated strain profiles at 0.5cm radius intervals from the center revealed the shape of the stress gradients in both conditions when facing the target (0^O^ Threshold Strain Horizon Angle; Figure 7C). We found that the threshold strain horizon angle was greater in the unstacked masses compared to the stacked masses (Figure 7D). In order to dissect the relative impacts of strain magnitude and area of strain distribution, we simulated strain fields in two conditions with mass distributions of comparable magnitude but different angles (Figure 7E,H) in the strain distributions shown (Figure 7F,G). Condition A contains 2 discs distributed so as to make a strain field the same approximate size as the 3 discs across the arena. In this case, the simulations result in similar threshold strain horizon angles for each side (Figure 7E,F,H), and experimental results confirm this prediction, with the *Physarum* choosing no side the vast majority of the time (Figure 7I). Thus, a similarly sized threshold strain horizon angle was sufficient to overcome a greater overall mass and prevented the *Physarum* from choosing either side. Condition B contains 3 discs on each side, with discs on one side distributed slightly more than on the other to increase the Threshold Strain Horizon Angle. In this case, the simulations result in a slightly larger threshold strain horizon angle on the side with the distributed discs (Figure 7G,H), and experimentally, a concomitant increase in choice of the “separated high mass” condition (Figure 7J), indicating that the slightly larger angle, even when the masses were exactly the same, increased the salience of the stimulus. These simulations and experiments support the argument that the sensed angle width of strain is critical to the decisionmaking process and suggesting that *Physarum* navigates toward specific masses by sensing the fraction of its perimeter impacted by the strain of a mass rather than the absolute magnitude of strain.

**Figure 7.**
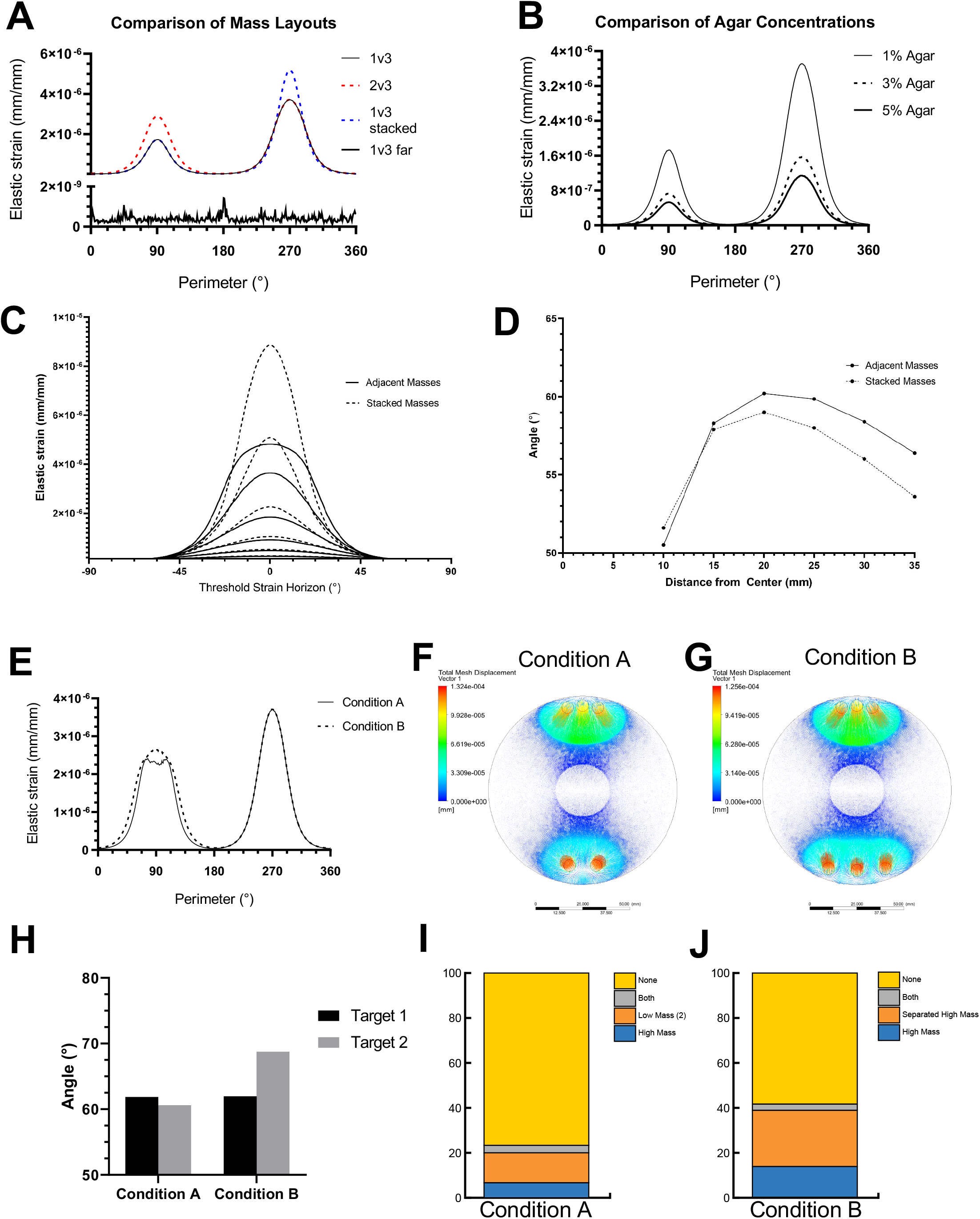
Width of threshold strain rather than magnitude strain explains mechanosensory decision making. (A) Simulated strain in 1%, 3%, and 5% agar gels at a radius of 30mm from the center shows both a decreased magnitude and angle width of strain propagation with increased agar concentration. (B) Simulated strain at a radius of 30mm for experimental conditions in 1% agar. (C) Simulated strain comparing adjacent and stacked 3-disc masses at 5mm radius intervals shows the differential magnitude and width of perceived strain oriented toward the high mass targets (0^O^). (D) The width of the strain above a threshold of 1.49×10^-7^mm/mm indicates a wider perceived angle for adjacent masses compared to stacked masses, supporting the hypothesis that the width of strain above a sensory threshold rather than the magnitude of strain drives *Physarum* mechanosensory decision making. (E) Plot of strain for two simulated conditions with similar magnitude strains but differential threshold strain angles. (F) Simulated strain distribution of Condition A. (G) Simulated strain distribution of Condition B. (H) Plot of threshold strain angles. (I) In the Condition A assay, the majority of the Physarum made no decision, suggesting that it did not find any difference between the two sides of the arena despite the 0.5mg weight differential because the strain angle was similar. () In the Condition B assay, 25% of the trials resulted in the Physarum choosing the condition where the discs were separated over just 14% where the Physarum chose the narrower angle despite the weight being equal.

### Theoretical model of *Physarum polycephalum* strain sensing

*Physarum* is widely known to grow in a pulsatile manner, which consists of a forward growth phase and a reverse streaming phase during which the cytoplasm is retracted away from the growth regions (Figure 8A). We have observed that this oscillation is critical for *Physarum* mechanosensation and that interrupting it by changing the substrate stiffness (Figure 4) or interfering with the oscillations (Figure 6) prevents accurate decision making. Thus, we propose a theoretical model of *Physarum* navigation where this oscillatory behavior acts as a sample-and-integrate function (Figure 8B): the growth regions sample the environment during the growth phase, optimize the direction of the network tubes during the reverse streaming phase by inducing internal tension in the *Physarum* network which then aligns future growth of the growth regions.

Finite element simulation indicates that *Physarum* navigates toward specific masses by sensing the fraction of its perimeter impacted by the strain of a mass rather than the absolute magnitude of strain. We hypothesize that *Physarum* is able to sense this strain gradient via establishment of a wider anchoring area to the substrate, providing a greater holding force during the reverse streaming phase of growth when the tube network is under tension. Less-anchored growth regions would therefore be more likely to retract slightly with each pulse, aligning sections of *Physarum* to the eigenvector of the strain vectors at each adhesion point. This proposed ratchet-like optimization mechanism between local adhesion sites could lead to long-distance network optimization. As prior studies have shown*, Physarum* optimizes the paths between locations/nodes by increased streaming in the network tubes that should also experience greater tension during reverse streaming. Our model also explains our observation that growth rates tend to increase as *Physarum* grows nearer a mass target where it experiences greater absolute and differential strain gradients. The summary of predictions and experimental data is presented in Table 2.

**Figure 8.**
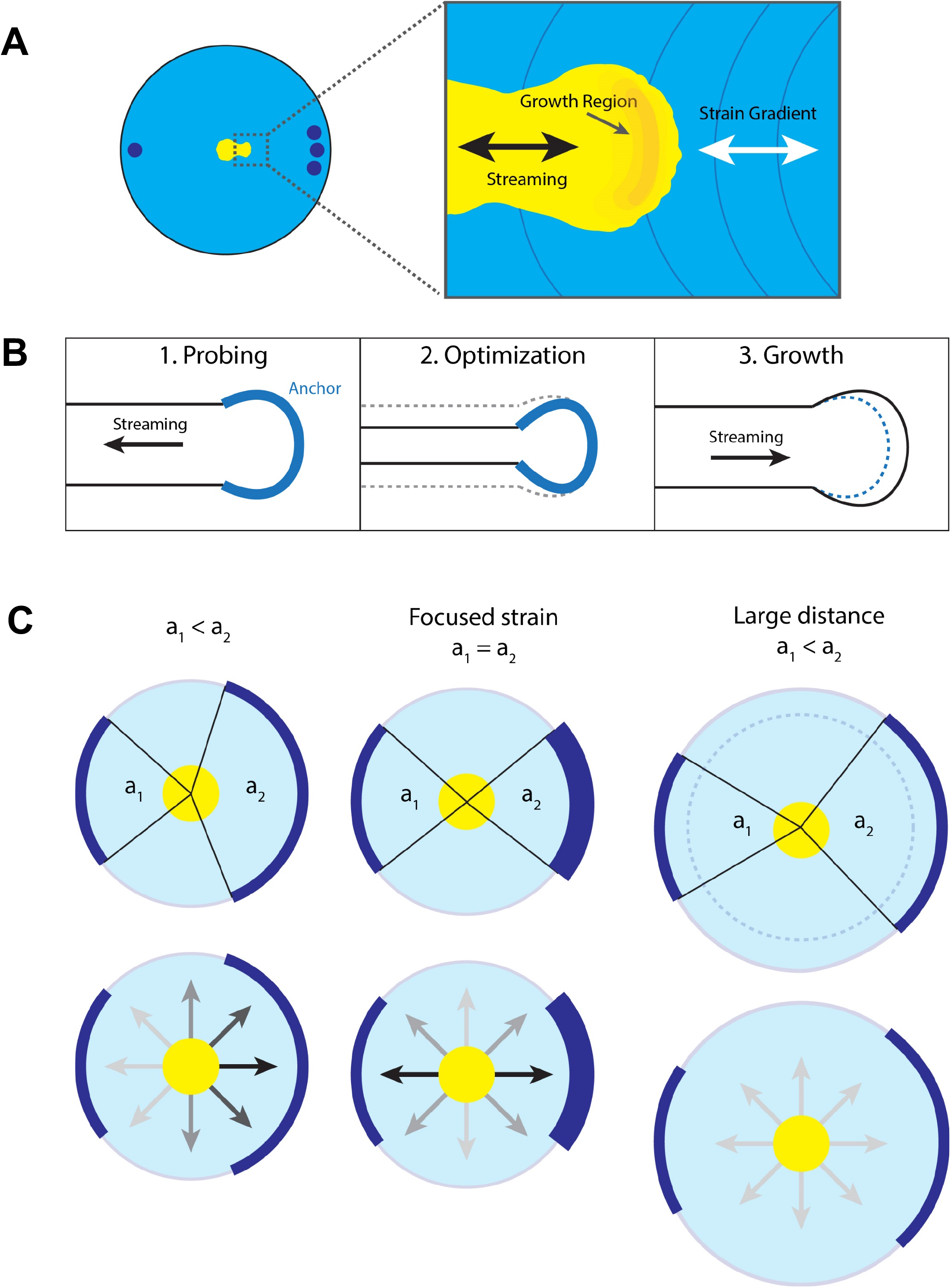
Proposed fluidically-coupled clutch model of mechanosensory navigation in *Physarum*. (A) Schematic of *Physarum* in a mass discrimination test. The inset illustrates the growth region along a strain gradient coupled with fluidic streaming. (B) Streaming integrates multiple strain-sensing growth regions where 1) focal adhesions anchor to the substrate and reverse streaming induces internal stress resulting in amplification of higher-stress focal adhesions and weakening lower stressed focal adhesions, thus acting as a rudimentary integrator or Eigenvector calculator to 2) incrementally optimize the *Physarum* network structure, followed by 3) forward streaming and subsequent incremental growth preferentially along stronger anchored projections. (C) Schematic of how angles of threshold strain rather than magnitudes are perceived by *Physarum* (top) and result in the observed decision-making probabilities (bottom).

**Table 2:** Computational model and its predictions matching to experimental data

| Experimental data shows that: |  |  | Claim | Does the theoretical model predict this? |
| --- | --- | --- | --- | --- |
| Static (No Shake) | Base Task | 1% agar 10cm dish<br>Decision to make: 3 vs 1 disc<br>Outcome: 3 discs | When distributed mass differential is high, growth tends toward the greater mass | Yes |
|  | Low Mass Differential | 1% agar 10cm dish<br>Decision to make: 3 vs 2 disc<br>Outcome: 3 discs | When distributed mass differential is low, growth tends toward the greater mass | Yes |
|  | Stacked Mass | 1% agar 10cm dish<br>Decision to make: 3 (stacked) vs 1 disc<br>Outcome: No significant differences between options | No discrimination is possible between a stacked high mass region and a low mass region | Yes |
|  | Large Distance | 1% agar 25cm dish<br>Decision to make: 3 vs 1 disc<br>Outcome: No Sig. Diff | Increasing the decision area between high and low mass regions inhibits discrimination | Yes |
|  | Mass Dispersion | 1% agar 10cm dish<br>Decision to make: 3 (dispersed) vs 1 disc<br>Outcome: Both | Dispersed high mass regions and low mass regions are equally selected | Yes |
|  | Proposal 2 | 1% agar 10cm dish<br>Decision to make: 3 vs 2 dispersed disc<br>Outcome: None | Increasing the dispersion of the low mass region does not change the preference towards high mass | Yes |
|  | Proposal 3 | 1% agar 10cm dish<br>Decision to make: 3 vs 3 dispersed disc<br>Outcome: None | Increasing dispersion while keep mass constant between choices, reveals an indecision between choices | Yes |
| Dynamic (Shake) | Base Task | 1% agar 10cm dish<br>Decision to make: 3 vs 1 disc<br>Outcome: Both | Mechanical disruption prevents discrimination between high and low mass regions regardless of spatial distribution | N/A |
|  | Low Mass Differential | 1% agar 10cm dish<br>Decision to make: 3 vs 1 disc<br>Outcome: Both |  | N/A |
|  | Stacked Mass | 1% agar 10cm dish<br>Decision to make: 3 vs 2 disc<br>Outcome: Both |  | N/A |
|  | Large Distance | 1% agar 25cm dish<br>Decision to make: 3 (stacked) vs 1 disc<br>Outcome: No Sig Diff. |  | N/A |
|  | Mass Dispersion | 1% agar 10cm dish<br>Decision to make: 3 (dispersed) vs 1 disc<br>Outcome: Both |  | N/A |
| Changing Shaking Frequency | 100% | 1% agar 10cm dish<br>Decision to make: 3 vs 1 disc<br>Outcome: Both | As the frequency of the mechanical disruption increases, the discrimination potential between masses decreases | N/A |
|  | 75% | 1% agar 10cm dish<br>Decision to make: 3 vs 1 disc<br>Outcome: Both |  | N/A |
|  | 25% | 1% agar 10cm dish<br>Decision to make: 3 vs 1 disc<br>Outcome: 3 discs |  | N/A |
| Anisotropic Gels | Parallel | 1% agar 10cm dish<br>Decision to make: 3 vs 1 disc<br>Outcome: 3 discs | Disrupting the strain fields orthogonal to the decision plane prevents high-low mass discrimination | Yes |
|  | Orthogonal | 1% agar 10cm dish<br>Decision to make: 3 vs 1 disc<br>Outcome: No sig. Diff. |  | Yes |
| Gel Density | 1% | 1% agar 10cm dish<br>Decision to make: 3 vs 1 disc<br>Outcome: 3 discs | As substrate stiffness increases, high-low mass discrimination decreases | Yes |
|  | 3% | 3% agar 10cm dish<br>Decision to make: 3 vs 1 disc<br>Outcome: No Sig Diff. |  | Yes |
|  | 5% | 5% agar 10cm dish<br>Decision to make: 3 vs 1 disc<br>Outcome: None |  | Yes |
| TRP<br>Channel<br>Inhibitor | GsTMX-4<br>(blocker) | 1% agar 10cm dish<br>Decision to make: 3 vs 1<br>disc<br>Outcome: Both | Chemically blocking TRP<br>channels inhibit high-low<br>mass discrimination | N/A |
|  | Linearized<br>peptide<br>(not blocker) | 1% agar 10cm dish<br>Decision to make: 3 vs 1<br>disc<br>Outcome: 3 discs | Sequence-matched,<br>linearized protein (negative<br>control) does not inhibit<br>high-low mass<br>discrimination | N/A |

This qualitative model explains our experimental data. Critically, it explains why the stacked mass condition resulted in equal probability of growth direction, unlike the clear directional growth in the adjacent mass condition and the lack of growth in the more distant placement of masses. Figure 8C summarizes the strain angles and magnitudes (top row) and directionality of growth (bottom row) that our model predicts and is observed experimentally. Although the absolute strain magnitude was comparable in all 3 cases, the stacked mass condition exerts the strain in a narrower angle than the condition with adjacent masses, leading to the lack of distinction between a single mass and three stacked masses and corresponding undirected growth (the “both” outcome).

## Discussion

### Aneural *Physarum polycephalum* can discriminate between masses at long range

Sensing and measurement of environmental conditions, as inputs to decision-making, are an essential capacity of living things, as relevant for unicellular organisms as for metazoan body cells during morphogenesis, regeneration, and cancer. To understand multicellularity, the origin of body architectures, and the rise of behavioral capacities throughout the tree of life, it is essential to characterize the ways in which evolution exploits physical forces for adaptive function. We developed a decision-making paradigm that identified a novel preference and sensing capability in the unicellular but large organism, *Physarum polycephalum*. A unique feature of this organism is that its behavior and its morphogenetic remodeling are the same process, providing a tractable window on the evolutionary scaling transition from cell behavior during pattern control to animal behavior during problem-solving (Fields et al., 2019; Keijzer et al., 2013; Keijzer, 2017).

*Physarum* reliably displayed a tropism for inert objects (glass disks) upon an agar surface arena, choosing to explore toward them even when no chemical signal (nutritional attractants) were present. Remarkably, it showed a reliable preference for objects with greater mass when presented with multiple options, demonstrating the ability to detect and compare the physical properties of aspects of its environment, and to actively grow out toward the preferred mass configuration. Quantitative morphological analysis and an unbiased machine-learning approach revealed that this capability included significant distance in both time and space: small pieces of *Physarum* were able to detect the presence of objects at a distance of several centimeters, and decide which way to go hours before actually moving in that direction. This reveals a minimum bound on the spatio-temporal boundary of the basal cognition of this organism (Levin, 2019).

### The *Physarum* body is a dynamic biomechanical sensory system

Distributing the mass or increasing the distance between the *Physarum* and its targets past a threshold was seen to interfere with mass preference. Similarly, when plates containing *Physarum* were exposed to repeated tilting or mechanical noise, frequency-dependent decrements of high-mass preference were observed. This suggested a biomechanical process. To further confirm our hypothesis and gain additional insight into how mechanosensation may allow mass-based decision-making at a distance, we used anisotropic agar gels. Establishing differential regions of distinct stiffness along or across the path between *Physarum* and the objects clearly demonstrated the importance of physical substrate properties to the ability of *Physarum* to perform mass sensing. We also identified a molecular component of this process. When treated with a known inhibitor of a mechanosensitive TRP channels, but not with a very similar control peptide, *Physarum* lost its discriminatory capacity.

We propose a model in which strain gradients, influenced by mass distribution across the substrate, provide the *Physarum* with an information-rich pathway through which to direct its growth. The *Physarum’s* body, extended across the agar gel, was able to probe the environment, optimize its distributed form, and grow along the strain gradient toward the target region. Suspecting shuttle streaming may be intrinsically linked to mechanosensory processing in the *Physarum*, we dynamically distorted gels with pulse patterns to match the organism’s intrinsic periodicities. We found that when a substrate-distorting piston matched the flow rate of the *Physarum*, the dynamic region was preferred over the static region (no piston). Thus, the *Physarum* can be fooled by a sort of “optical illusion” in the mechanical sensing space of this organism, mimicking with artificial vibrations the kind of signal that the slime mold’s sonar-like pulsations would normally receive from an attractive environmental mass.

### Computational modeling of mass sensing

We formulated a quantitative model of *Physarum* mechanosensory information processing which builds on the clutch model of large-scale tissue mechanosensing proposed for mammalian tissues (Sunyer et al., 2016), exploiting highly-conserved basic principles of biomechanics. Crucially, this model predicted the specific outcomes which we observed (Table 2). Due to the long, non-linear spans between growth regions in *Physarum* we propose that fluid flow and resulting oscillations in cellular tensioning drive the information integration process instead of direct cytoskeletal coupling. Evolutionarily speaking, this proposed mechanism is intermediate between the cytoskeleton-mediated mechanosensing of mammalian cells and cell wall-reliant mechanosensing of bacteria and plants, particularly related to osmolarity stress and loss of turgor pressure. In fact, fungal hyphae have been observed to grow in an oscillatory manner and bidirectionally transmit signals tied to pathogen attack and nutrient sources (Schmieder et al., 2019). *Physarum* may have optimized this inherent ability to execute information processing more rapidly.

### Evolutionary perspective: biomechanical sensing from cells to organisms

Mechanosensing is an evolutionarily conserved sensory modality present in all living organisms, from prokaryotes (Belas, 2014) to fungi (Kumamoto, 2008), plants (Monshausen and Gilroy, 2009), and animals (Ingber, 2006; Luo et al., 2013), as a way of processing environmental cues to promote survival. Mechanical cues have been demonstrated to directly induce specific biological programs, including switching between growth and apoptosis (Chen et al., 1997). In eukaryotes, this process is mediated by a suite of mechanosensitive ion channels and cellular adhesion proteins that connect cells to their microenvironment at focal adhesions (Arnadottir and Chalfie, 2010; Ingber, 2006; Luo et al., 2013). The cytoskeleton plays a critical role, particularly in mammalian cells, in transmitting mechanical cues among focal adhesions and the mechanosensitive ion channels in the plasma membrane (Ingber, 2006, 2008). Importantly, mechanical forces continue to be exploited when unicellular organisms merge into multicellular bodies, and multicellular clusters transmit mechanical information among the constituent cells, which leads to more accurate and sensitive mechanosensing (Sunyer et al., 2016). Biomechanics, and the use of physical forces to sense the environment, is now known to be crucial in embryogenesis (Campas et al., 2014; Davidson, 2012; Nelson, 2009), stem cell biology (Mammoto et al., 2011), and cancer (Discher et al., 2005; Oudin and Weaver, 2016), driving the need for new model systems in which to understand how biological tissue exploits physical forces to make decisions about growth and form.

Advanced environmental sensing and decision-making capacities of complex plants and animals evolved from cell-level capacities (Baluška and Levin, 2016; Lyon, 2006). While it’s not known why *Physarum* prefers to explore large masses over small ones, future work on *Physarum* in its native environment is likely to reveal the adaptive advantages of prioritizing the exploration of larger masses. Overall, this capacity reveals a novel, evolutionarily ancient nexus in which physical forces have been exploited for the simultaneous control of body morphogenesis and problem-solving behavior. Evolution discovered the utility of exploiting physical phenomena, such as bioelectricity (Levin and Martyniuk, 2018) and biomechanics (Beloussov, 2012), very early. However, cells did not lose these capacities when joining to form multicellular collectives, because development and maintenance of complex bodies requires cells to sense and make decisions based on information from a much larger size scale. Thus, biomechanical mechanisms have been widely utilized across metazoan, including roles in osteogenesis/remodeling, neurogenesis, immune etc. (Darnell et al., 2017; Pageon et al., 2018). Our discovery of mechanosensation in the unicellular slime mold underscores the early evolutionary origin of this computational modality (Cox et al., 2018). Interestingly, it is a truly multi-scale phenomenon, working not only at the level of single cells but also exploited by entire organisms, such as for example spiders that decode the vibrations in their webs to identify location of prey (Mortimer et al., 2019), analogously to the abilities of *Physarum* within its substrate.

### Conclusion

Future work in this fascinating field will extend the model toward molecular-biological investigation of complex decision-making, and the biophysical processes by which sensory information is integrated, stored, and processed toward subsequent remodeling and behavior. We are especially focused on using electrophysiological and other imaging techniques to learn to read the “engram” – the internal biophysical states by which *Physarum* represents its mapping of the external environment and the results of its sensing activities. An understanding of this phenomenon in simpler models may advance the decoding efforts in neuroscience, which also seeks to understand how behavioral information is stored in living tissue. There are perhaps scale-invariant and substrate-indpendent principles of learning and memory that extend to all cognitive system that can be deduced using this simple model organism (Baluška and Levin, 2016; Friston et al., 2015; Pezzulo and Levin, 2015; Vallverdu et al., 2018; Yokawa and Baluška, 2018). Moreover, measurement of physical properties of the environment is an important but poorly-understood aspect of metazoan morphogenesis, including for example the biomedically important feedback loops that allow cells in a regenerating organ or remodeling structure to determine how to grow and when to stop based on large-scale properties of the real-time anatomy (Butler et al., 2009; Levin and Martinez Arias, 2019; McLaughlin and Levin, 2018; Whited and Levin, 2019). Thus, well beyond contributions to fundamental knowledge in evolution, cell biology, and basal cognition, the understanding of physical forces as an information medium for cell behavior and spatial control is likely to have important implications for regenerative medicine and perhaps serve as biological inspiration for artificial living machines and soft robotics (Doursat and Sanchez, 2014; Doursat et al., 2013; Kamm and Bashir, 2014; Kamm et al., 2018; Kim et al., 2013; Rieffel et al., 2014).

## Acknowledgments

We thank Dr. Audrey Dussutour for providing the sclerotic *Physarum polycephalum*, Jayati Mandal for general laboratory assistance, and members of the Levin Lab for their helpful comments and discussions. We thank Dr. Nicolas Rouleau for his insightful comments and statistical suggestions. We gratefully acknowledge Dr. Joshua Finkelstein for his support and helpful discussions. This research was supported by the Allen Discovery Center program through The Paul G. Allen Frontiers Group (12171), and this research was also sponsored by the Defense Advanced Research Projects Agency (DARPA) under Cooperative Agreement Number HR0011-18-2-0022, the Lifelong Learning Machines program from DARPA/MTO. The content of the information does not necessarily reflect the position or the policy of the Government, and no official endorsement should be inferred. Approved for public release; distribution is unlimited.

## Methods

### Culturing conditions

*Physarum polycephalum* sclerotia, the encysted resting state of Australian origin (provided by the Dussutour Lab, Toulouse, France), were re-hydrated, split, dehydrated on filter paper, and stored in darkened conditions at 20 °C. Each sclerotic body was reconstituted two weeks prior to the experimental treatment by moistening filter paper containing dehydrated *P. polycephalum* with sterile water. For experiments, we used the hydrated, plasmodial stage of *P. polycephalum*. The plasmodia were cultured on 150mm cell culture dishes covered in a layer 25mm thick of 1% w/v agar (Fisher Scientific, USA). Cultures were maintained in a temperature-and humidity-controlled (90% humidity and 22 °C) incubator in the dark. Flakes of rolled oats (Quaker Oats, USA) were liberally spread over the surface of the agar to provide a nutrient-rich environment to promote growth and expansion. The cultures of plasmodia were sub-cultured onto new plates every two days.

### Decision-making assay

We developed an *in vitro* assay using various masses and distributions of inert glass microfiber discs (GE Healthcare, Life Sciences, USA) to assess the decision-making power of *Physarum polycephalum*. The inert glass microfiber discs were positioned on the agar at the periphery of petri dishes as indicated in each experiment. The discs did not contain any nutritional value – when the nutrient source (oats) was replaced with discs of equal weight, *Physarum* displayed a starvation response that confirmed the disc-only condition was equivalent to nutrient deprivation.

Discrimination tasks were administered within the context of an arena which consisted of either a 10cm diameter Petri dish (or a 25cm diameter Petri dish for the “large distance” conditions) filled with 1% non-nutrient agar at a thickness of 0.5cm. Plasmodial blocks of 1cm diameter were resected from the sub-cultures and placed into the center of individual arenas. Each sample fragment always contained 1 oat flake to ensure the presence of a nutrient source and eliminate the contribution of hunger. Glass microfiber discs were placed equidistant and in opposite direction to the *Physarum* (Fig 1A). Two mass-related variables were experimentally manipulated: the number of discs positioned at each extremity of the plasmodia (i.e., one, two, or three) and the orientation of the discs (i.e., stacked or side-by-side). If placed side-by-side, discs were oriented orthogonal to the axis that runs through the two-disc loci and the plasmodium and were separated by 4.5mm (10.5mm for “large distance” conditions). Thus, the plasmodia were exposed to combinations of binary choices (e.g., three stacked discs versus one disc; three side-by-side discs versus one disc).

The arenas were then placed in a darkened incubator for 24 hours, after which time decisions were recorded by photographing each plate using a Canon EOS Rebel T7i DSLR camera and positioned over a light pad (Artograph, USA). Decisions were quantified as the plasmodium travelling ≥75% of the linear distance between its point of origin and the center of the disc region. Four possible decision outcomes were identified: 1) high mass, 2) low mass, 3) both, or 4) none. “High-mass” refers to growth toward a three-disc region (stacked or side-by-side), “low mass” refers to growth toward a one-disc region, “both” refers to ≥75% distance travelled in both directions, and “none” refers to conditions where distance thresholds were not met (<75% distance travelled) in either direction. Choices, plate orientation and location within the incubator (e.g., shelf height) were counterbalanced to eliminate confounding factors.

To create time-lapse recordings, arenas were placed within darkened acrylic boxes outfitted with a flatbed scanner controlled by VueScan software. Images were captured every 15 minutes over a 24-hour period to measure incremental changes in orienting behavior as a function of the experimental conditions.

### Mechanical Disruption

We developed a mechanical perturbation system to observe whether exogenous disruption can interfere with the decision-making process. Instead of placing arenas on immobile, level shelves within the incubators during the growth period (as in the baseline decision making assays), each assay was placed inside an incubator on a laboratory-grade tabletop rocker, and the *Physarum* was allowed to make the decision on this surface with constant rocking (Benchmark Scientific Inc., USA). The tilt of the table was set to displace each arena 22.5° above and below the central plane with a periodicity of 0.75Hz over the 24-hour growth period (Fig. 3A). The axis of tilt was across the disc-plasmodium-disc midline. The maximum output of the tilt table was reduced to examine the step-wise effects of increased motion on plasmodial selection. To isolate the contribution of continuous movement from any effects due to tilt itself, some plates were grown on static tilt tables that were locked into their downward-tilted conformation (22.5° below the central plane). The arena was oriented on the platform such that the three-disc or one-disc region was placed at the lowest-most point on the tilt table platform. Some plates were continuously tilted (0.75Hz); however, the axis of tilt was orthogonal to the disc-plasmodium-disc axis.

### Mechanosensitive Channel Inhibition

The water soluble, stretch-activated ion channel inhibitor, GsMTx-4 (Abcam, USA)(Gnanasambandam et al., 2017) was used to block mechanosensitive channels in the *Physarum*. The decision-making assay was repeated as described above with the addition of 60μL (to a final concentration of 30μM) of the prepared drug, which was placed on the central mass of the plasmodial fragment at the center of the arena upon plating. *P. polycephalum’s* outermost shell is highly porous, making the organism relatively permeable to water-soluble drugs. Our observations indicated that full absorbance was achieved within 20 minutes of depositing the drug. Once the drug absorption had been observed, the plates were placed in the incubator for the 24-hour growth period before measurement.

### Anisotropy Assay

A 3D-printed petri dish insert (Fig 4A) was made using poly lactic acid (PLA, description data file available as Supplemental Figure 5). The file was uploaded and printed by a 3D printer (Fortus 360mc from Stratasys) at the Bray Lab of Tufts University, producing an insert made of ABS-M30. The insert consisted of longitudinal strips of PLA interrupted by longitudinal spaces (void). The insert was submerged in 1% agar in a petri dish to generate a pattern of PLA-agar-PLA-agar which disrupted the isotropic substratum. A thin strip of agar was positioned over PLA strips to achieve a surface uniformity. The net result was an anisotropic arena consisting of polar-oriented strips of intermittent high-and low-density substrata on which plasmodia were placed (see Fig 4B) The disc-plasmodium-disc axis of the dish could be positioned in along the longitudinal axis of the anisotropic substrate or orthogonal to it. In this way, we could assess whether access to a longitudinal strip of agar or PLA would promote orienting-behavior to enhance decision-making capacities.

### Finite Element Modeling

3D modeling of patterned agar substrates was performed using computer-aided design (CAD) software (Dassault Systèmes SolidWorks, Waltham, MA). Finite element simulation of mechanical behavior was performed using the mechanical structural module of the Ansys software package (Ansys, Canonsburg, PA). The goal of the simulation was to estimate the strain magnitudes and directions caused by the presence of masses on the surface of the model to analyze experimental conditions. The interaction of the masses on the surface of the agar was simulated by applying gravity to the model. Since the growth of *Physarum* occurs only on the surface of the 6.35mm thick agar substrate, a high-resolution mesh was applied to the agar surface and surrounding applied to achieve more accurate visualization of the strain and stress magnitude distribution. The material properties used in the model are shown in Table 1.

### Dynamic Compression Device

We used a Dual Mode Muscle Lever (Aurora Scientific, Model # 300C) to provide pressure stimuli on the agarose medium to disrupt pulsing-mediated mechanosensing. The 300C system has 2 operating modes, a Force mode and a Length Mode, thus called Dual Mode. For our experiments we used the Force mode which allowed the system to convert voltages into force. Given the specified 3üüC’s conversion rate of 100 micro-Newton/Volt, we sent periodic amounts of voltages to the 300C system to generate small oscillatory forces on the agarose medium. The 300C system is programmatically driven by our customized Matlab GUI and a National Instruments USB6002 (DAQ) device. To ensure that the gel is not punctured by the lever arm, we replaced the 3üüC’s lever arm with blunt cap with diameter of 1cm, the same diameter of the static discs

### Data acquisition and statistical analysis

Images obtained after the 24-hour growth period were processed in ImageJ. Linear distance between the central point of the arena (origin of the plasmodia) and the disc region was measured. A threshold at seventy five percent of the plate were used to determine decisionmaking on each side. Categorical data were entered into SPSS v20. The frequency of each decision option was computed and converted to a percentage that reflected the relative tendency to select a given option when presented with two targets. Weighted chi-squared tests were performed to determine whether there was a significant difference between the expected and observed frequencies of the decision categories.

### Dynamical machine learning model

Videos of *Physarum* growth in the decision-making arena were collected (N=43). Frames collected at 15-minute intervals from videos of *Physarum* completing the decision-making assay (Supplemental Fig. 3A) were processed. First, images were rotated and cropped, leaving out the discs so that the model was only learning from the features of the growth and not the environment. Then, growth area masks were automatically extracted from the image of the plate the *Physarum* covered. Finally, growth area increment masks were created from the difference between the previous image and the current image (Supplemental Fig. 3B).

In each frame, the machine learning task was to predict the target direction of the growth (right or left) considering the observed growth dynamics. As a machine learning algorithm, we used Gradient Boosted Trees (GBT, XGBoost library realization (Chen, 2016)) with cross-entropy loss. In this study, GBT operates on a vector of hand-crafted features extracted from the images, including: normalized barycenter of a recent 15-minute *Physarum* outgrowth increment as well as its leftmost and rightmost relative positions, recently covered and cumulative areas of growth, and the compressed history of these features weighted by the Fibonacci sequence from the start of growth to the current frame. See Supplemental Fig. 3D for the model input and output specification, Supplemental Fig. 3C for an approximate visualisation of the features, and Supplemental Fig 2E for the predictions given by the model for the samples from Supplemental Fig 3A-C.

We validated the proposed method using the leave-one-out cross-validation (LOOCV) approach: in each iteration during LOOCV, we fit a model to N–1=42 videos and evaluate it on the single, remaining one. Although this process is computationally expensive, it is useful for small datasets like we have (Hawkins et al., 2003). During the training and testing of the model, the images were flipped in every possible way to remove the bias.

To compare our statistical model with a coin-toss, weakly stationary process with ½ mean and 1/(4*N*) variance we used OLS regression combined with an Augmented Dickey-Fuller (ADF) test (Said, 1984) for trend stationarity to exclude a possible unit root.

## Supplemental Materials

**Supplemental Figure 1.** Video of *Physarum polycephalum* tugging on substrate.

**Supplemental Figure 2. Video of *Physarum* outgrowth.** Video of the time lapse image depicted in Figure 1B. *Physarum* (yellow) grows in the 100mm arena placed in the dark. Images were captured every 15 minutes for 24 hours.

**Supplemental Figure 3. Dynamical machine-learning model predicting *Physarum* growth.** (A) Time-lapse videos of growth in the 1 vs. 3 assay used to develop a machine learning model that attempts to predict the future growth direction from the morphological structure of the *Physarum*, thus enabling detection of the time at which it has made a decision about the mass distribution in its environment. (B) Automatically generated incremental growth masks. (C) Graphical representation of the features used by the machine learning algorithm at each time frame for the prediction of the target direction (darker color shows older outgrowth). (D) Prediction results for the data shown on Fig. A-C. (E) Model’s input-output specification. (F) Prediction accuracy as a function of time: the light blue line corresponds to the mean accuracy for the prediction; the thicker blue line corresponds to the trend in the accuracy improvement; thin dashed blue lines correspond to the 95% confidence interval for the measured accuracy; the dashed red line corresponds to the naive deterministic classifier that outputs the destination that the growth is closest to at the moment. (G) Temporal dynamics of the growth: solid lines correspond to the median distances from the growth extremities (nearest — blue, furthest — red) to the target; shaded areas correspond to the second and third quartiles of the population. (H) Trends in the most recent growth directions: the solid line corresponds to the median distance from the recent outgrowth center of mass to the target; shaded areas correspond to the second and third quartiles of the population. (F-H) The vertical dashed line indicates the time at which the prediction accuracy becomes statistically significantly higher than random; at this point, the majority of the *Physarum* samples still grow in an opposite direction to the target.

**Supplemental Figure 4.** Baseline assay for *Physarum* growth in a field. *Physarum* was placed in an arena with no disks and assessed for frequency with which it grew towards specific, arbitrarily defined regions of the arena. The values identified here are used as the “expected” values for χ-squared analysis.

**Supplemental Figure 5.** CAD files for anisotropic gel mold. The STL/SLDPRT file contains a file format for the anisotropic mold to be uploaded and printed by a 3D printer to produce an insert made out of ABS-M30. This mold is then submerged in agar in a petri dish to generate the anisotropic substrate.

